# Uncertainty aversion predicts the neural expansion of semantic representations

**DOI:** 10.1101/2023.01.13.523818

**Authors:** Marc-Lluís Vives, Daantje de Bruin, Jeroen M. van Baar, Oriel FeldmanHall, Apoorva Bhandari

## Abstract

Correctly identifying the meaning of a stimulus requires activating the appropriate semantic representation among many alternatives. One way to reduce this uncertainty is to differentiate semantic representations from each other, thereby expanding the semantic space. In four experiments, we test this semantic-expansion hypothesis, finding that uncertainty averse individuals exhibit increasingly differentiated and separated semantic representations. This effect is mirrored at the neural level, where uncertainty aversion predicts greater distances between activity patterns in the left inferior frontal gyrus when reading words, and enhanced sensitivity to the semantic ambiguity of these words in the ventromedial prefrontal cortex. Two direct tests of the behavioral consequences of semantic-expansion further reveal that uncertainty averse individuals exhibit reduced semantic interference and poorer generalization. Together, these findings demonstrate that the internal structure of our semantic representations is shaped in a principled manner: aversion to uncertainty acts as an organizing principle to make the world more identifiable.

## Introduction

Human life is rife with uncertainty, from the information we gather through our senses^1,2^ to the unpredictable outcomes of our actions^3–6^. People intolerant to uncertainty find it especially aversive and are therefore strongly motivated to reduce it^7,8^. When confronted with ambiguous situations that lack a clear interpretation, those who are averse to uncertainty experience stress and will take action to avoid any additional uncertainty^9^. Because these individuals are more sensitive to uncertainty and perceive greater uncertainty than those who are uncertainty tolerant^10^, they are often better at remembering concepts or cues that signal uncertainty^11^. In short, uncertainty aversion plays an outsized role in shaping behavior across a range of domains^12–15^. Despite this, uncertainty is inescapable; it even imbues the concepts we use to make sense of the world^16^. Take for instance the words “slip” and “lapse”, which have similar, but subtly different meanings. When focusing on the similarities, both refer to some form of mistake, which means that they can represented similarly in semantic space and it may not matter which word one uses. Focusing on the differences, however, highlights that a “slip” is usually trivial and accidental, while a “lapse” can be serious and imply responsibility, and this may cause an individual to separate these concepts in semantic space. Here, we propose that individuals seeking to minimize uncertainty distinguish concepts from one another by separating them in semantic space.

Semantic representations encode our conceptual knowledge about the world and influence how we perceive incoming sensory information^17^, which enables us to ascribe meaning to stimuli^16^. However, because stimuli typically activate more than one concept, there is always some uncertainty when identifying meaning. The degree of uncertainty between a pair of concepts depends on their separation in semantic representational space. Therefore, ensuring that concepts are sufficiently differentiated from one another in such a space helps to reduce semantic uncertainty. This logic accords with classic work demonstrating that the closer a pair of concepts are in psychological space (e.g., snow and cold), the more confusable they are, while more dissimilar concepts are easier to discriminate and thus suffer far less from semantic interference (e.g., cat and black)^18,19^. In short, the distances between concepts in semantic representational space determine whether one’s everyday experiences are conceptually ambiguous or fairly clear-cut. We hypothesize that individuals averse to uncertainty mitigate the effects of uncertainty by making their semantic representations more distinct. Here we test whether this process, which we term *semantic expansion*, causes increases in pair-wise distances between concepts in psychological or neural space, thus rendering related concepts as more separable for a downstream readout mechanism.

The echo of such a strategy should also be detectable in the structure of neural representations. Consider a brain region that encodes concepts in the activity of its neurons. Each concept is associated with a distinct neural activity pattern, and separating the activity patterns from each other as much as possible enables a downstream neuron reading out the representation to disambiguate each concept from the others^20–22^. The greater the distance between neural activity patterns, the more separable the representation is^23,24^, and, consequently, the less uncertainty there is in the representation—which would be very useful to those who prefer to reduce the attendant uncertainty of a concept (Fig. 1). Simply put, our semantic expansion hypothesis predicts that individuals who are averse to uncertainty should exhibit more differentiated neural activity patterns.

**Figure 1.**
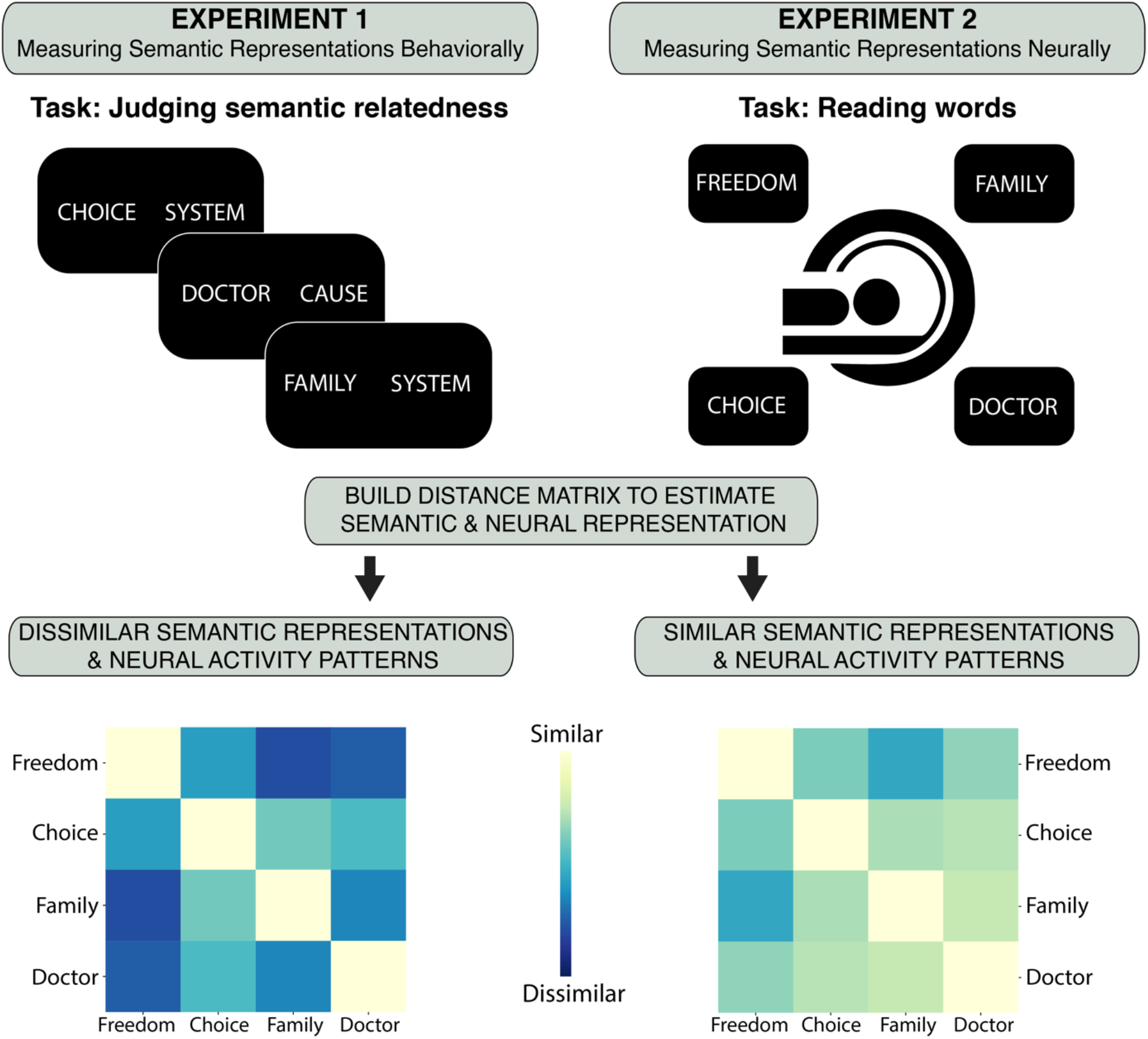
Dissimilar semantic representations and neural activity patterns are more identifiable. Each word is associated with a semantic representation captured by judging the semantic relationship between two sets of words (Experiment 1) or a unique neural pattern of activity captured by reading a list of words one at a time (Experiment 2). Semantic and neural competition increases as similarity between neural activity increases. Neural representations that are more dissimilar on average would suffer less neural competition and, as a consequence, be more identifiable.

While the need to discriminate between concepts encourages concepts to be separated from each other, there is an opposing functional influence on the structure of the representational space: the need to keep similar concepts closer to each other in psychological space. This promotes useful generalization between similar concepts. In other words, the structure of a semantic representation reflects a trade-off between the need for discrimination and generalization. We hypothesize that people averse to uncertainty, through semantic expansion, privilege discrimination over generalization.

To test this semantic-expansion hypothesis, we conducted four experiments where we examined the effects of individual differences in uncertainty attitudes on semantic distances between words in psychological and neural space and their behavioral consequences. Results reveal that uncertainty aversion promotes the separation of semantic representations at both the psychological (Experiment 1) and neural levels (Experiment 2). We then tested the direct behavioral consequences of the semantic-expansion by examining how uncertainty aversion shapes the trade-off between discriminability and generalization. We find that uncertainty aversion reduces semantic interference (Experiment 3) and leads to poorer generalization (Experiment 4). In other words, people averse to uncertainty exhibit more distinct semantic representations, which results in them prioritizing the need to discriminate between concepts, at the cost of their ability to generalize.

## Results

### Aversion to uncertainty is associated with more distinct semantic representations

In Experiment 1, 103 participants made semantic relatedness judgements between 16 target words and 42 comparison words (see Methods and Fig.1). Similarity between the 16 target words was then estimated by correlating the 42-element vectors of relatedness ratings across words. To calculate distances between words using the same number of meaningful dimensions for each participant^25^, multidimensional scaling (MDS) was applied. A four-dimensional partition was selected given that it was the lowest number of dimensions that produced a good fit (stress = 0.09)^26^. In this four-dimensional space, the average distance between words was computed for each participant and then correlated with individuals’ uncertainty attitudes, which were captured by the well-validated intolerance of uncertainty scale (IUS). The IUS assesses uncertainty aversion by asking people to rate to what extent statements like “the ambiguities in life stress me” describe them^9^. Demonstrating a semantic expansion effect, we found greater aversion to uncertainty was associated with an increase in the distance between concepts in psychological space (r = 0.35, p = 0.0003; Fig. 2A). This relationship was not dependent on the number of dimensions selected when applying multidimensional scaling since the same result also holds for three, five, and six dimensions (see Supplementary Table 1). This effect was not driven by an increase in response variability in people with higher IUS scores: there was no significant relationship between response variability and uncertainty aversion (r = -0.03, p-value = 0.79, see Supplementary Figure 1), and the semantic expansion effect remained significant after controlling for response variability in a regression (= 0.001 ± 0.0002 (S.E.), p < 0.001). The effect also remains significant after controlling for global variables like IQ, age, and gender (β = 0.001 ± 0.0003 (S.E.), p = 0.002, see Supplementary Table 2).

**Figure 2.**
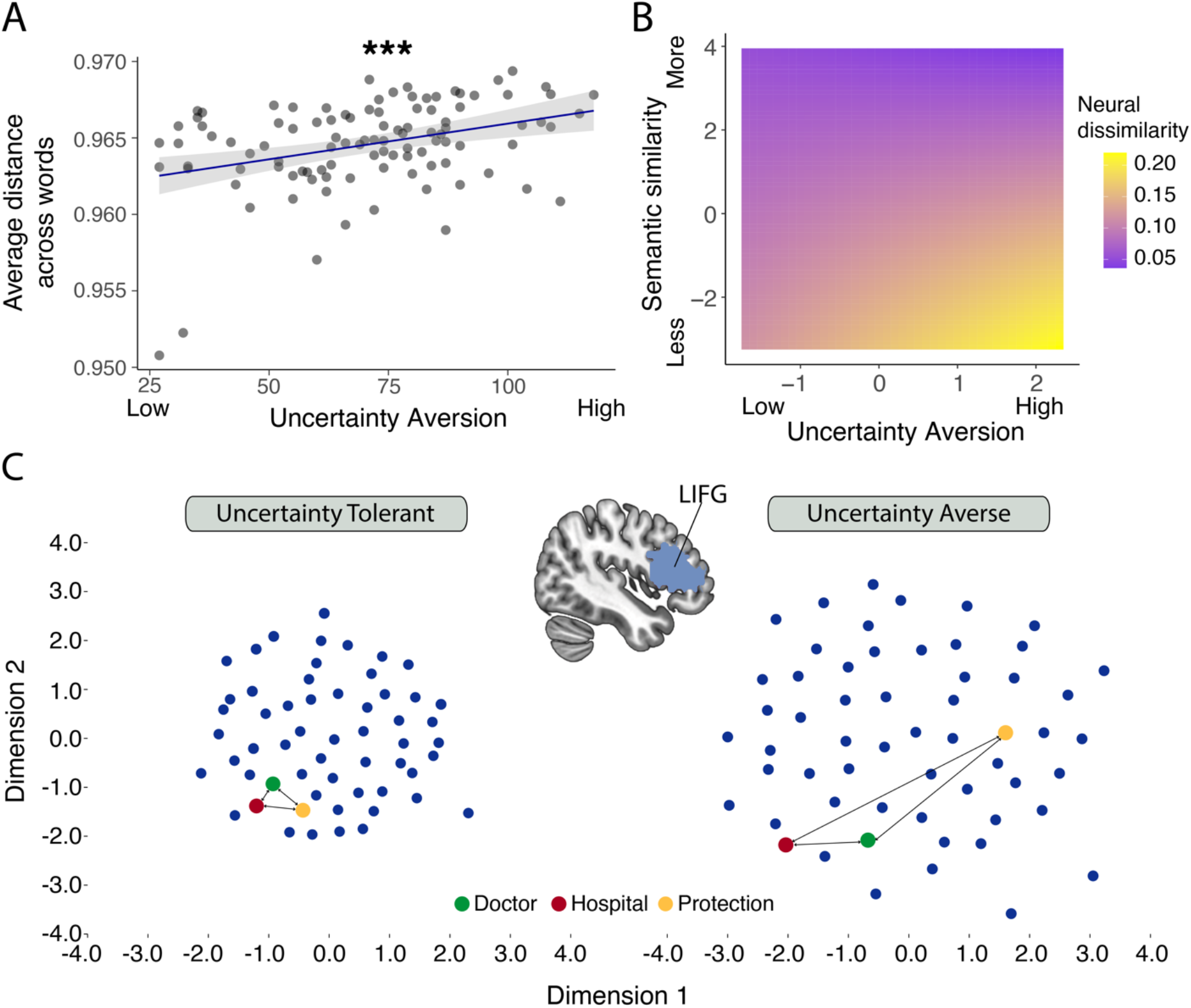
**A**. A significant correlation was observed between uncertainty intolerance and distances between semantic representations. Shaded area reflects 95% confidence intervals, the result remains significant even after removing the two outliers. *** p < 0.001. **B**. Uncertainty aversion modulates the relationship between semantic similarity and neural similarity. Semantic representations were captured by a reduced neural space for people tolerant of uncertainty, while the neural space was larger for people averse to uncertainty. Cross-validated Mahalanobis distances between pairs of words plotted as a function of beta estimates extracted from the regression model. **C**. This relationship between uncertainty aversion and representational distance between words in the LIFG is further illustrated using two representative participants that differed in their uncertainty attitudes. Multidimensional scaling was applied to their average neural RDMs to transform the neural representational distances for all word pairs into a two-dimensional space. Each word is represented by a point in the plane. Words are more distant from one another for the uncertainty averse participant.

### Semantic neural representations expand as a function of the aversion to uncertainty

To test whether uncertainty aversion is also associated with more distal neural representations, in Experiment 2 we analyzed data from a functional magnetic resonance imaging (fMRI) experiment in which 44 participants read 60 words and were asked to think about their meaning (see Methods). We used representational similarity analysis (RSA) to estimate the distance between words in neural semantic space. Given previous work showing a role for the left inferior frontal gyrus (LIFG), angular gyrus, middle temporal gyrus, anterior temporal cortex, and perirhinal cortex in semantic representation^27–32^, we hypothesized that the pair-wise distances between the neural representations of words in these brain regions (localized to the left hemisphere^33^) would be sensitive to uncertainty aversion. We further reasoned that the ventromedial prefrontal cortex (VMPFC), as well as the precuneus—regions known to process various forms of uncertainty^6,34–38^, and in some cases have also been implicated in semantic processing (i.e., the precuneus)^31^—might be involved in indexing the relationship between uncertainty intolerance and how words are represented.

In any neural semantic representation, the neural pattern distance between a pair of concepts should be proportional to their semantic dissimilarity. We reasoned that, under the semantic expansion hypothesis, uncertainty aversion would increase the slope of the relationship between semantic dissimilarity and neural representational dissimilarity, as people averse to uncertainty expand their neural semantic space. To test this prediction, for each participant, a neural representational dissimilarity matrix (RDM) was constructed for each ROI by computing the cross-validated Mahalanobis distance between all word pairs^39^. A model RDM was also constructed containing the pairwise differences of the semantic dissimilarity between words derived from Global Vectors for Word Representation (GloVe; see Methods). For each of our regions of interest, a linear mixed-effect regression was run with the vectorized lower triangle of the neural RDM as the dependent variable, and the vectorized lower triangle of the semantic similarity RDM, an individual’s tolerance to uncertainty, and their interaction, all as predictors. In the LIFG, we observed a significant positive interaction between semantic similarity and aversion to uncertainty: increasingly dissimilar words exhibit increasingly dissimilar activity patterns, especially for individuals who are averse to uncertainty (β = 0.02 ± 0.006 (S.E.), p = 0.005; Bonferroni-corrected for multiple comparisons across seven ROIs, Fig. 2C). Including age, gender and level of education as covariates did not change this result (see Supplementary Results Experiment 2). This effect was not observed in any other ROIs (see Supplementary Table 3).

We also tested the possibility that this neural semantic expansion effect was driven by increased executive control exerted by participants with high IUS scores, which could in theory produce an online reshaping of the semantic representation. To test this, we estimated the mean activity in the fronto-parietal regions of the brain^40^, which are known to index executive control and mental effort^40^, and included them as covariates in our regression. Again, the significant interaction between semantic similarity and IUS remained significant (β = 0.03 ± 0.005, p < 0.001).

### Aversion to uncertainty is associated with greater neural sensitivity to semantic ambiguity

The uncertainty associated with multiple semantic representations cued by a given word is higher for words that have multiple meanings and are thus considered semantically ambiguous. For example, ‘run’ can be used in a diverse array of contexts whereas ‘artichoke’ is typically only used in the context of food. A word that can be used in many different contexts has a larger number of referents and greater semantic ambiguity^41^. For those intolerant to uncertainty, encoding this ambiguity would enable semantic representations to be tagged as ones that must be structured more distally. To test this, we used RSA to estimate the pair-wise neural pattern distances between each word. Given that each word is associated with a certain degree of semantic ambiguity—estimated using a corpus that counts the number of contexts a given word appears in^41^—we can interrogate whether our regions of interests^6,34,35,37,38,42^ track this semantic ambiguity as a function of an individual’s aversion to uncertainty. We constructed a neural RDM comprised of the pair-wise cross-validated Mahalanobis distances between individual words, and a model RDM containing the pairwise differences of the semantic ambiguity between words derived from a language corpus (see Methods). We then ran a linear mixed-effects regression analysis where the vectorized neural RDM served as the dependent variable, and the vectorized semantic ambiguity RDM and an individual’s tolerance to uncertainty (as well as their interaction) served as predictors for our regions of interest.

Given the importance of semantic ambiguity to uncertainty-averse individuals, we expected ambiguity to be encoded more strongly for participants with higher IUS. Indeed, in the VMPFC, we observed a significant positive interaction between semantic ambiguity and aversion to uncertainty: words that are similar in their semantic ambiguity exhibit similar activity patterns, especially for individuals who are averse to uncertainty (*β* = 0.03 ± 0.006 (S.E.), *p* < 0.001; Fig. 3B). A similar effect, albeit of lower magnitude, was also observed in the precuneus and LIFG, as indexed by a significant interaction between intolerance of uncertainty and semantic ambiguity (precuneus: *β* = 0.02 ± 0.006 (S.E.), *p* = 0.03, LIFG: *β* = 0.02 ± 0.006 (S.E.), *p* = 0.04—these effects, however, do not remain significant after controlling for increased cognitive effort presumably associated with processing semantically ambiguous words (see Supplementary Table 4 for the rest of the ROIs).

**Figure 3.**
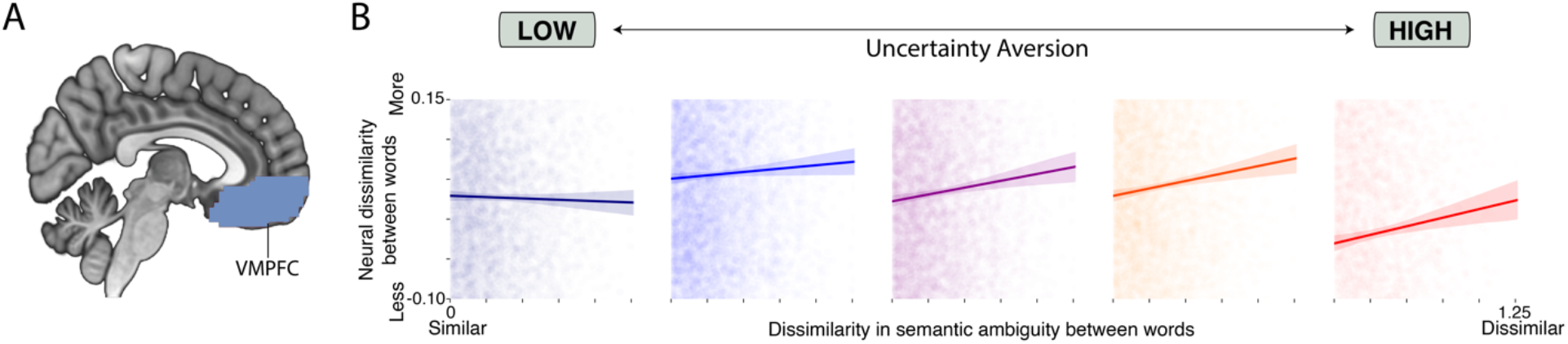
**A**. Ventromedial prefrontal cortex (VMPFC) was a pre-defined region of interest. **B**. For visualization purposes, participants were binned into 5 groups based on discrete IUS intervals. For each participant, a neural RDM was estimated, vectorized, and then correlated with the vectorized semantic ambiguity RDM derived from the language corpus. For those averse to uncertainty, activity patterns in the VMPFC are more dissimilar between words that are semantically ambiguous. Shaded areas reflect 95% confidence intervals.

### Aversion to uncertainty improves semantic discrimination

The semantic-expansion hypothesis posits that uncertainty-averse subjects reduce uncertainty by increasing the separation between concepts in both psychological and neural space. A natural consequence of this hypothesized semantic expansion is that for those intolerant to uncertainty, similar concepts can be more easily discriminated and are less likely to produce semantic interference. To test for this reduction in semantic interference, we leveraged the classic Deese-Roediger-McDermott (DRM) false memory paradigm where subjects first encode lists of words and are then asked to discriminate between old and new words in a surprise recognition memory test (Fig. 4A). A robustly observed effect within the literature is that lure words semantically similar to the initial words (e.g., snow and cold) tend to be falsely endorsed as old^15,16^.

**Figure 4.**
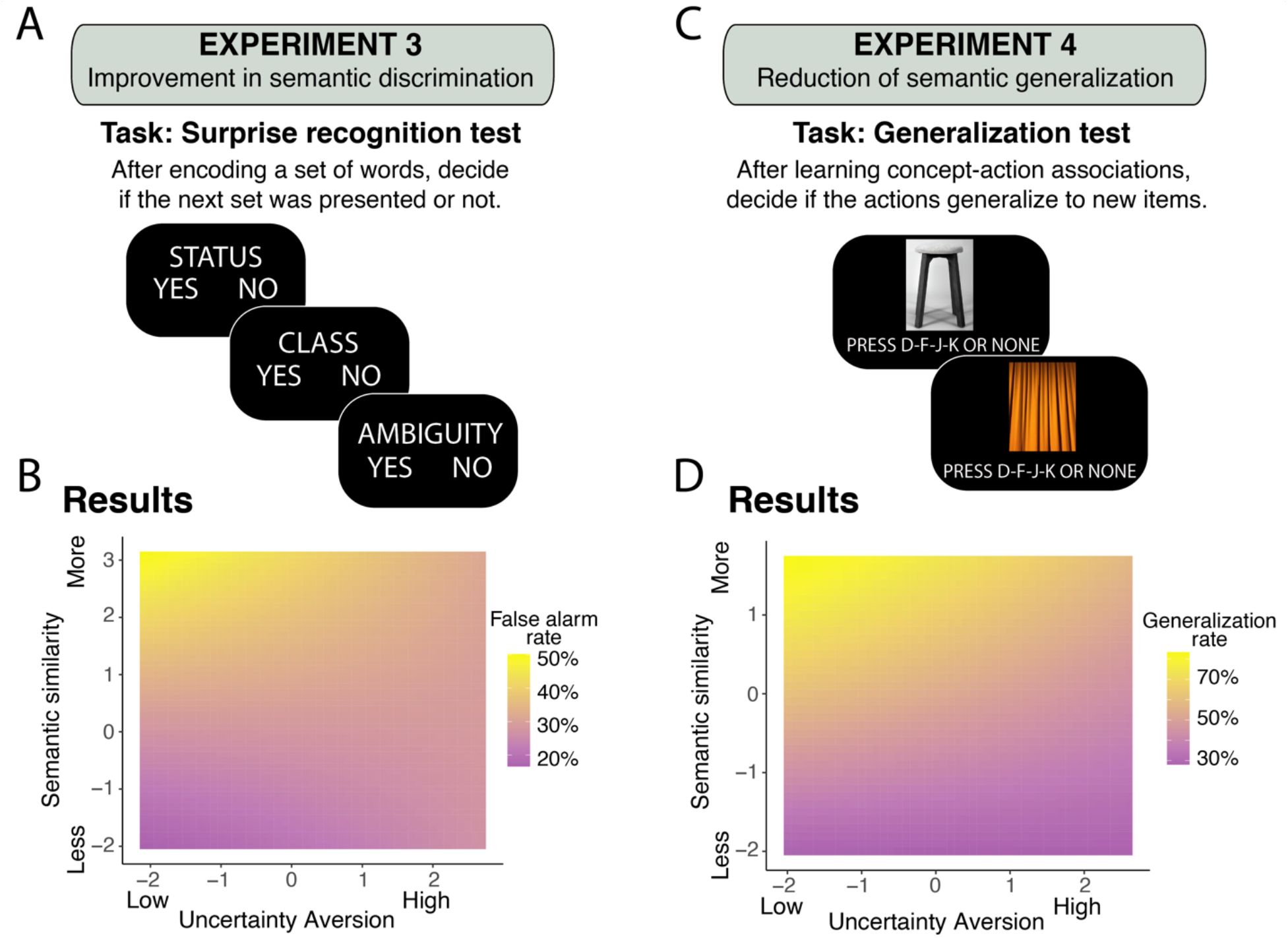
**A**. A surprise recognition test was administered to participants to test for an increase in semantic discrimination for people averse to uncertainty. **B**. Uncertainty aversion modulates the effect of semantic interference during a memory recognition test. While those who can tolerate uncertainty do falsely report recognizing words that are similar to previously presented words, semantic interference is significantly lower for those who are averse to uncertainty. **C**. A generalization task was administered to test for a decrease in semantic generalization for people averse to uncertainty, **D**. Uncertainty aversion modulates the effect of semantic similarity during a generalization task. Semantically closer pictures supported stronger generalization, but this effect was attenuated in those averse to uncertainty. False alarm rates (B) and generalization rates (D) plotted as a function of beta estimates extracted from respective regression models.

In Experiment 3, 206 participants first studied 50 words, and were then given a surprise recognition memory task, with 100 words, 50 of which were the ones previously shown, and 50 of which were novel (see Methods and Fig. 4A). Participants performed the task reasonably well, correctly identifying the old words 72% of the time (hit rate) while misidentifying the new words 31% of the time (false alarm rate). To test the prediction that uncertainty aversion would modulate the effect of semantic interference, we focused our analysis on the false alarm rate. We first estimated semantic similarity between the novel and previously seen words using an unsupervised learning algorithm trained on a large language corpus (GloVe). The maximum similarity between previously seen words and each new word was calculated and used as a regressor (alongside participants’ IUS scores, as well as their interaction) to predict the false alarm rate. Results reveal that as the similarity between novel and previously seen words increases, so too does the probability of falsely identifying a novel word as being already seen (β = -0.23 ± 0.11 (S.E.), p = 0.03)—an effect that was modulated by one’s tolerance to uncertainty (interaction: β = 0.06 ± 0.02 (S.E.), p = 0.01, Fig. 4B). That is, the greater the aversion to uncertainty is, the better the discrimination between similar concepts. This effect remains significant when adding average hit-rate as a regressor in the model (β = 0.06 ± 0.02 (S.E.), p = 0.01, see Supplementary Figure 2).

### Aversion to uncertainty reduces semantic generalization

Classic work shows that generalization between stimuli decreases as the psychological distance (i.e., discriminability) between the stimuli increases^43^. By this logic, a direct consequence of semantic expansion—which is observed in those averse to uncertainty—should be a reduction in semantic generalization. To test this, in Experiment 4, 197 participants first learned an association between four concepts (e.g., chair, wrench, which was cued by images of objects) and four actions (pressing one of four possible keys). To learn the associations, participants were presented with an image of an object and told to press one of the keys. Feedback about whether the correct key was pressed was immediately provided. Once they learned which concept was associated with which key, they moved to a generalization phase where they were presented with new pictures of other concepts (e.g., stool, screwdriver). This time, they were instructed to press whichever key they saw fit (no feedback was provided during the generalization phase, see Methods and Fig. 4C). For each of the four concepts presented during learning, two concepts were selected as stimuli in the generalization phase, one which was semantically close to the initial concept (e.g., stool is conceptually similar to chair) and another that was further away (e.g., curtain is conceptually distant to chair), but still semantically closer than any of the other target concepts (e.g., curtain is even more distant to jellyfish than chair). Pressing the same key associated with the concept during learning was coded as successful generalization (=1), while any other response was coded as lack of generalization (=0). Semantic similarity between the novel concepts during generalization and initial concepts during learning was estimated using GloVe to predict the probability of generalization, alongside participant’s aversion to uncertainty and their interaction. As expected, results reveal that the probability of generalization increased as the similarity between novel and target concepts increased (*β* = -0.83 ± 0.042 (S.E.), *p* < 0.001), an effect that was modulated by uncertainty aversion (interaction: *β* = 0.08 ± 0.03 (S.E.), *p* = 0.01), such that semantically similar concepts supported a lower level of generalization as uncertainty aversion increased (see Fig. 4D).

## Discussion

Uncertainty is present in virtually all aspects of our lives, from perceptual judgments to decision outcomes. A major source of uncertainty comes from deciphering meaning in a conceptually ambiguous world. Not only are most stimuli associated with more than one meaning^44^, but a stimulus can also activate multiple similar semantic representations^45^ (e.g., slip and lapse), which results in greater uncertainty. Here, we directly tested one strategy for reducing this type of uncertainty, which is to expand one’s semantic representations.

Behavioral and neural results illustrate the semantic-expansion hypothesis. Aversion to uncertainty scales with more differentiated semantic representations (Experiment 1), which is also associated with increasingly separated neural activity patterns in the LIFG (Experiment 2)—a region classically involved with matching words with their appropriate semantic representation^46–49^. Two direct behavioral tests of this semantic-expansion hypothesis demonstrate that uncertainty averse individuals exhibit improved semantic discrimination during a surprise memory recognition test (Experiment 3) and reduced semantic generalization for similar concepts (Experiment 4). As uncertainty becomes more aversive, the structure of semantic representations becomes increasingly separated so that each concept is more easily identified, and as a consequence, the mapping between stimulus and meaning becomes more certain. Together, these findings demonstrate that personality needs, such as the need to reduce uncertainty, can shape fundamental aspects of cognition, including how we represent meaning. Indeed, our results suggest that an organizing principle of semantic representations might be the amount of uncertainty people can tolerate.

The fact that we found that the expansion of neural activity patterns was localized to the LIFG— and not a broader neural network associated with uncertainty^50–52^—suggests that regions not traditionally related to uncertainty processing also play a distinct role in helping to reduce semantic uncertainty. Taken a step further, rather than relying on the arbitration of specific neural regions to encode various types of uncertainty^50–52^, it is possible that the brain reduces uncertainty by shaping the geometry of neural activity patterns associated with concepts (or stimuli). This account dovetails with current, broader theories of neural activity that posit a relationship between distinct representations and reduced task interference^20,23,24^.

Our findings also demonstrate that classic uncertainty regions, namely the VMPFC and precuneus, are especially sensitive to semantic uncertainty in those averse to uncertainty—the first work to link the neural encoding of semantic representation to regions involved in processing uncertainty^27^. Considering that the semantic expansion result was localized to the LIFG, this finding suggests a neural division of labor. In a situation devoid of context in which virtually any meaning associated with each word could be retrieved, VMPFC appears to signal the differences in the number of possible referents that could be retrieved. In contrast, the LIFG plays a role in biasing the structure of semantic representations—through the alteration of the geometry of neural representations—which expands depending on one’s aversion to uncertainty, thus avoiding semantic confusion. This accords with prior work linking LIFG activity with the resolution of semantic ambiguity^53^.

Even though the specific role of the left IFG in language processing is still debated^54^, it has been implicated in semantic control^55,56^. The fact that the observed semantic expansion effect was localized only to the LIFG may suggest that semantic expansion does not occur in stable semantic representations, but is instead produced online based on task demands. We caution, however, against interpreting the LIFG as the sole player involved in processing semantic expansion for two reasons. First, the fact that we did not observe semantic expansion in the anterior temporal cortex may simply be because our study was not optimized to capture BOLD signal in this region. Second, the online account is hard to reconcile with our finding in Experiment 3 that semantic interference is reduced in those averse to uncertainty (i.e. since there is no task demand to discriminate concepts during encoding when interference occurs, but only later, during a surprise recognition memory test). Indeed, similar semantic interference effects have previously been associated with medial temporal regions^18^ rather than LIFG. Future work should clarify whether semantic expansion is a property of the stored representation, the result of an online computation, or both.

Regardless, the resolution of semantic ambiguity through semantic expansion has behavioral consequences that can be understood through the lens of a computational tradeoff between discrimination and generalization^23,43,57^. The world would be impossible to navigate if we had to learn from scratch how to engage with each new stimulus. Instead, we routinely generalize what we learn about from one stimulus to other conceptually similar stimuli^43,57,58^. This sensitivity to the similarity between concepts, however, comes at the cost of increased uncertainty and the possibility of interference. We are more likely to confuse two similar concepts with one another, resulting in errors of discrimination. A natural consequence of semantic expansion is that it shifts this tradeoff in favor of discrimination. Increasing the separation of concepts in a psychological and neural semantic space would make concepts more discriminable. We show that people averse to uncertainty are indeed less affected by interference between similar semantic representations and show improved discrimination between concepts. Simultaneously, they are also less able to generalize behaviors between similar concepts. In other words, uncertainty aversion leads to people making their world more compartmentalized, trading off generalization in favor of better discriminability and lower uncertainty.

Important open questions remain. First, does semantic expansion preferentially shape concepts that are closer together, or generally influence the whole representation? When the structure of semantic representation was measured (Experiments 1 and 2), we found no direct evidence that the expansion effect is greater for similar concepts. On the other hand, in Experiments 3 and 4, people averse to uncertainty exhibited greater semantic discrimination (Experiment 3) and poorer generalization (Experiment 4), especially for very similar concepts. It is possible that the different task demands in these latter experiments made our measurements more sensitive for detecting the semantic expansion effect with concepts that share increasing similarity, and future work can help arbitrate whether this is the case. Second, might our findings be explained by expanding the dimensionality of the representation? Previous theoretical work has argued that the dimensionality of a representation controls a tradeoff between discriminability and generalizability^23^, and indeed, an increase in dimensionality would result in greater separability. In line with this view, we found that those averse to uncertainty had higher discriminability (and thus lower uncertainty) at the cost of poorer generalizability (Experiments 3 and 4). However, we found no direct evidence that uncertainty aversion is associated with higher dimensionality (Experiments 1 and 2, see Supplementary Figures 3 & 4). It is possible that, given the low dimensionality of the representational structures that we examined, and the poor sensitivity of current dimensionality measures, our analysis was not sufficiently powered to detect higher dimensionality for people averse to uncertainty, which leaves it as an open question for future work.

Research has investigated how diverse experiences alter mental and neural representations, mostly by measuring different levels of expertise (e.g., trained musicians vs. non-musicians^59^). Rather than showing how different experiences shape neural representations, the semantic-expansion hypothesis proposes that differences can originate even when people share the same experience. Based on the notion that those averse to uncertainty are more likely to adopt strategies that decrease uncertainty and increase identifiability, we have demonstrated that this stable personality trait predicts a more distal structure of semantic representations—a mechanism that could be applied across domains (e.g., animals, objects) and cognitive processes (e.g., perception).

## Methods

All experiments were approved by Local University’s Institutional Review Board.

### Experiment 1

#### Participants

108 people participated in the experiment on Prolific. 5 participants were excluded after they failed to pass an attention check at the end of the experiment, resulting in a final sample of 103 participants (47 women, 51 men, 3 non-binary, and 2 transgender; mean age 34, SD = 12).

#### Word Stimuli

For consistency, we selected a subset of words that were also used in Experiment 2, which were originally selected to address an orthogonal research question. Apart from these words, we identified other words that the target words could be compared with. To do this, we searched through lists of abstract words and selected 48 words that we i) classified as abstract (e.g., life, system), ii) were comprehensible, and iii) varied in how similar they were to each other. A table with these words can be found in the supplementary material with their summary characteristics.

#### Procedure

Participants completed 672 relatedness ratings between 16 target words and 42 comparison words on a scale from 1 (not related) to 9 (very related). The target and comparison words were presented on the screen simultaneously and participants then judged to what extent they were related. The instructions did not ask subjects to focus on a specific semantic domain. Words were selected to be relatively abstract: freedom, family, choice (see full list in the Supplementary Material). Following this task, participants’ uncertainty attitudes were captured using the 27-item Intolerance of Uncertainty scale as well as demographic information. The experiment lasted around 40 minutes and participants received $4.50 as compensation.

### Experiment 2

#### Participants

44 participants, all of whom were right-handed (17 women and 27 men and; mean age 32, SD = 14) participated in the experiment. The data analyzed here was collected as a part of a larger study exploring the neural mechanisms of intolerance to uncertainty. Participants provided informed consent and received $40 as monetary compensation.

#### Procedure

Participants completed a battery of tasks inside and outside the MRI scanner. Before the MRI scanning session began, participants provided basic demographic information and completed a list of questionnaires including the 27-item Intolerance of Uncertainty scale^9^. The scanning session lasted approximately 1.5 hours, in which they completed a Word Reading Task and a Video Watching Task (not analyzed in this research). After the scanning session, participants completed other questionnaires and tasks that were also not analyzed for this research.

#### Word Reading Task

The word reading task was conducted inside the MRI scanner. The task consisted of 6 runs, each made up of 80 trials. In every run, 60 unique words were individually presented on the center of the screen without any accompanying context. Each trial stimulus was presented for 2.5s with a fixed inter stimulus interval of 2.5s during which only a fixation cross appeared on the screen. 20 null-trials were added in which only a fixation cross was shown to improve the efficiency of the design for estimating stimulus evoked activity to each word. The trial order was randomized, and the words were presented in black font on a white background screen. To maintain participant’s attention, participants had to indicate if they believed the word presented was political or non-political by pressing one of two response buttons. Total run duration was approximately 6 minutes and each participant completed 6 runs.

#### fMRI acquisition and preprocessing

MR images were collected on a Siemens Prisma Fit 3-Tesla research-dedicated scanner. T2*-weighted functional scans were acquired using a multi-slice sequence capturing three slices at once to ensure whole-brain coverage with short repetition time (TR = 1500 ms). 60 3-mm transverse slices were acquired, each with 64×64 voxels of 3.0 mm isotropic, building up a field of view (FOV) that covered the entire brain except part of the cerebellum. The FOV was tilted upward by 25 degrees at the front of the brain to minimize tissue gradient-related signal dropout in the orbitofrontal cortex. Contrast settings were optimized for cortical grey matter (TE = 30 ms, flip angle = 86°). T1-weighted anatomical scans were acquired using a standard MPRAGE sequence (160 sagittal slices with 256×256 voxels of 1.0 mm isotropic, TR = 1900 ms, TE = 3.02 ms, flip angle = 9°). Preprocessing was performed using *fMRIPrep* 1.5.1rc2^60^ (RRID:SCR_016216), which is based on *Nipype* 1.3.0-rc1^60,61^ (RRID:SCR_002502).

#### Anatomical data preprocessing

The T1-weighted (T1w) image was corrected for intensity non-uniformity (INU) with N4BiasFieldCorrection^62^, distributed with ANTs 2.2.0^63^ (RRID:SCR_004757), and used as T1w-reference throughout the workflow. The T1w-reference was then skull-stripped with a *Nipype* implementation of the antsBrainExtraction.sh workflow (from ANTs), using OASIS30ANTs as target template. Brain tissue segmentation of cerebrospinal fluid (CSF), white-matter (WM) and gray-matter (GM) was performed on the brain-extracted T1w using fast^64^ (FSL 5.0.9, RRID:SCR_002823). Volume-based spatial normalization to one standard space (MNI152NLin2009cAsym) was performed through nonlinear registration with antsRegistration (ANTs 2.2.0), using brain-extracted versions of both T1w reference and the T1w template. The following template was selected for spatial normalization: *ICBM 152 Nonlinear Asymmetrical template version 2009c* ^65^ (RRID:SCR_008796; TemplateFlow ID: MNI152NLin2009cAsym).

#### Functional data preprocessing

The following preprocessing was performed for each of the BOLD runs per participant. First, a reference volume and its skull-stripped version were generated using a custom methodology of *fMRIPrep*. The BOLD reference was then co-registered to the T1w reference using FLIRT^66^ (FSL 5.0.9) with the boundary-based registration^67^ cost-function. Co-registration was configured with nine degrees of freedom to account for distortions remaining in the BOLD reference. Head-motion parameters with respect to the BOLD reference (transformation matrices, and six corresponding rotation and translation parameters) are estimated before any spatiotemporal filtering using mcflirt^68^ (FSL 5.0.9). BOLD runs were slice-time corrected using 3dTshift from AFNI 20160207^69^ (RRID:SCR_005927). The BOLD time-series (including slice-timing correction when applied) were resampled onto their original native space by applying a single, composite transform to correct for head-motion and susceptibility distortions. These resampled BOLD time-series referred to *preprocessed BOLD*. The BOLD time-series were resampled into standard space, generating a *preprocessed BOLD run in MNI152NLin2009cAsym space*. Several confounding time-series were calculated based on the *preprocessed BOLD*: framewise displacement (FD), DVARS and three region-wise global signals. FD and DVARS are calculated for each functional run, both using their implementations in *Nipype*^70^. The three global signals are extracted within the CSF, the WM, and the whole-brain masks.

Additionally, a set of physiological regressors were extracted to allow for component-based noise correction (*CompCor*^71^). Principal components are estimated after high-pass filtering the preprocessed BOLD time-series (using a discrete cosine filter with 128s cut-off) for the two *CompCor* variants: temporal (tCompCor) and anatomical (aCompCor). tCompCor components are then calculated from the top 5% variable voxels within a mask covering the subcortical regions. This subcortical mask is obtained by heavily eroding the brain mask, which ensures it does not include cortical GM regions. For aCompCor, components are calculated within the intersection of the aforementioned mask and the union of CSF and WM masks calculated in T1w space, after their projection to the native space of each functional run (using the inverse BOLD-to-T1w transformation). Components are also calculated separately within the WM and CSF masks. For each CompCor decomposition, the k components with the largest singular values are retained, such that the retained components’ time series are sufficient to explain 50 percent of variance across the nuisance mask (CSF, WM, combined, or temporal)^72^. The remaining components are dropped from consideration.

The head-motion estimates calculated in the correction step were also placed within the corresponding confounds file. The confound time series derived from head motion estimates and global signals were expanded with the inclusion of temporal derivatives and quadratic terms for each^73^. Frames that exceeded a threshold of 1.0 mm FD were annotated as motion outliers. All resamplings can be performed with a single interpolation step by composing all the pertinent transformations (i.e., head-motion transform matrices and co-registrations to anatomical and output spaces). The data were not susceptibility distortion corrected in the absence of fieldmaps. The Gridded (volumetric) resamplings were performed using antsApplyTransforms (ANTs), configured with Lanczos interpolation to minimize the smoothing effects of other kernels^74^. Non-gridded (surface) resamplings were performed using mri_vol2surf (FreeSurfer).

#### fMRI data cleaning

For the word reading task, two exclusion criteria were used. First, we excluded runs in which there were excessive motion artifacts, defined as a framewise displacement greater than 1 mm in more than 10% of the repetition times. Second, runs in which participants did not press the response button because of experimenter error or a lack of attention were also excluded. This resulted in a total exclusion of 8 runs from 6 participants (on average 1.3 runs per participant and not more than 2 runs for each participant, which leaves at least 4 presentations of each word for each participant) and the exclusion of one participant all together, leading to a final inclusion of 43 participants for the statistical fMRI analyses.

#### Statistical fMRI analysis

##### Regions of Interest (ROI)

Based on previous research on semantic processing^27–32^, we defined our ROIs as the left inferior frontal gyrus (LIFG), angular gyrus (AG), middle temporal gyrus (MTG), anterior temporal cortex (ATC), and perihinal cortex. The LIFG was defined using the Triangular and Opercular part of the LIFG, as stipulated by the Automated Anatomical Labeling (AAL) atlas^75^. The AAL atlas was also used to demarcate the AG, MTG, and ATC. The perirhinal cortex was defined as Brodmann Area 35 and 36. We also had *a priori* ROIs related to uncertainty^6,34–38^, which included the ventromedial prefrontal cortex (VMPFC) and precuneus—which were also demarcated using the AAL.

##### Representational Similarity Analysis (RSA)

For every participant, a General Linear Model (GLM) was constructed using SPM12 in MatlabR2017b, in which every word that was presented in the word reading task was modelled as a separate regressor yoked the duration of stimulus display. Motion was regressed using 6 motion directions, their first derivatives, their squares, and the first derivatives of the squares. In addition, the CompCor physiological regressors described above were included as covariates for denoising. Fitting this model to the data yielded a beta map per run, per word. The beta values for each voxel were extracted for each ROI. The RSA Toolbox^76^ (http://github.com/rsagroup/rsatoolbox) was then used to compute, for each participant and ROI, a neural representation dissimilarity matrix (RDM) using the cross-validated Mahalanobis distance between all word pairs. This measure incorporates multivariate noise normalization and cross-validation, which leads to higher reliability of the neural distance estimates^39^.

##### Testing for semantic expansion

To investigate whether intolerance of uncertainty expands the neural representation of semantics, we adopted a linear mixed-effects model in which the neural representational dissimilarity of each word pair was regressed by uncertainty attitudes and semantic dissimilarity as captured by GloVe. A neural RDM by for each ROI was computed using the approach described above. A model RDM containing the pairwise dissimilarity in semantic dissimilarity was created by computing the difference between semantic similarity scores obtained from GloVe. The neural RDM was vectorized, z-scored, and included in the regression model. All others factors included in the regression model were also z-scored. The main effects of IUS and semantic dissimilarity and the interaction between the two was tested predicting the vectorized, scaled, neural dissimilarity patterns in the ROIs, with participant included as a random intercept^77^.

##### Semantic ambiguity analysis

To investigate whether intolerance of uncertainty modulates how semantic ambiguity is processed, we adopted a linear mixed-effects model in which the neural representational dissimilarity of each word pair was regressed by uncertainty attitudes and semantic ambiguity. A neural RDM for each ROI was computed using the approach described above. A model RDM containing the pairwise dissimilarity in semantic ambiguity was created by computing the difference between semantic ambiguity scores obtained from the English Lexicon Project Web Site^78^ (https://elexicon.wustl.edu/). The neural RDM was vectorized, z-scored, and included in the regression model. All others factors included in the regression model were also z-scored. The main effects of IUS and semantic ambiguity and the interaction between the two was tested predicting the vectorized, scaled, neural dissimilarity patterns in the ROIs, with participant included as a random intercept^77^.

### Experiment 3

#### Participants

213 participants were run on Prolific. 7 participants were removed since they failed to pass an attention check at the end of the experiment, and 1 participant because they did not complete the whole experiment, totaling in 205 participants (103 women, 96 men, 4 non-binary, and 2 gender fluid; mean age 29, SD = 6).

#### Words Stimuli

Concepts were selected to be relatively abstract and include a mix of relatively similar (e.g., status-class) and dissimilar (e.g., ambiguity-peculiarity) new and old words.

#### Procedure

Participants were first presented with a word and a fractal and were told that these two stimuli were associated. After this, participants were asked to judge the likelihood that 10 other words were also associated with the same fractal on a scale from 1 (not likely) to 7 (very likely). Participants completed 5 blocks of this task. At the beginning of each block, a new fractal was associated with a new word, and participants had to judge the association between the fractal and 10 new, comparison words. In total, they completed 50 trials of this task for a total presentation of 50 comparison words. A surprise recognition test was then administered. In total, participants were presented with 100 words (50 old, 50 foils, for a subset of participants, a coding error made the presented foils to be only 48), and had to indicate whether they had seen this word before (Yes or No). Finally, participants were asked to judge the similarity between the 50 word-pairs that they saw in the associative task on a scale from 1 (not similar) to 7 (very similar). Again, uncertainty attitudes were collected using the IUS scale together with demographic information. The experiment lasted around 15 minutes and participants received $2 as compensation.

#### Analysis procedure

A mixed-effect binomial regression was performed on the probability of falsely recognizing a lure (1=yes, 0=no) predicted by the maximum similarity of the old words for each lure, uncertainty attitudes, and the interaction between the two. Subject and word item were both added as independent random intercepts.

### Experiment 4

#### Participants

221 participants were run on Prolific. 13 participants were removed since they failed to pass an attention check at the end of the experiment, 9 participants were removed because their average accuracy in the learning phase was lower than 70%, and 2 participants were removed because their average reaction time in the generalization task was lower than 500 milliseconds, totaling in 197 participants (94 women, 97 men, 6 non-binary; mean age 29, SD = 6).

#### Procedure

The experiment consisted of two phases, a learning and a generalization phase. In the learning phase, participants learned the association between four pictures cueing different concepts (e.g., jellyfish, chair, canoe) and four keys (e.g., “d”, “f”, “j”, “k”). On each trial, a picture was presented and participants pressed one of the four keys. Immediately after, participants received feedback regarding whether the selected key was the one associated with the concept. Participants responded to 20 trials for each association-pair in a pseudo-randomized order. After learning, participants completed the generalization phase, in which new pictures of other concepts were presented and participants had to decide whether to select one of the old keys or the space bar that represented a “no response”. Two concepts were selected for each of the four target concepts, one that was very semantically close to the target (e.g., chair-stool), and another one that was semantically further (e.g., chair-curtain), for a total of eight pictures depicting eight different concepts. Each picture was presented twice in a pseudo-randomized order. In total, participants completed two independent blocks of learning and generalization phase for a total of 8 pair-associates (see Supplementary Material for the list of concepts presented). Presentation order was counterbalanced across participants. At the end of the experiment, uncertainty attitudes were collected using the IUS scale together with demographic information. The experiment lasted around 15 minutes and participants received $2 as compensation.

#### Analysis procedure

A mixed-effect binomial regression was performed on the probability of generalization (1=yes, 0=no) predicted by the similarity of the new concept with the target concept, uncertainty attitudes, and the interaction between the two. Subject and category were both added as independent random intercepts.

